# Morphological and yield responses of *Cannabis sp.* to night-time disruption

**DOI:** 10.1101/2023.12.28.573579

**Authors:** Bryan Steve González-Vanegas, Felipe Sarmiento Salazar, Camila Calderón, Valentina Osorio

**Affiliations:** Universidad Nacional de Colombia, sede Bogotá, Facultad de Ciencias Agrarias; Universidad Nacional de Colombia, sede Bogotá, Facultad de Ciencias; One Tropical Seeds S.A.S

**Keywords:** Cannabis sp., Night Break, flowering, photoperiodism, cannabinoids, physiology

## Abstract

The production of *Cannabis sp.* under inductive photoperiods requires investment in lighting infrastructure to delay the inflorescence stage and maintain yield. Previous reports have shown that the interruption of the night phase (Night Break, NB) prevents floral development, however, the effects related to stress response, yield and cannabinoid concentration are unknown for cannabis. The present study sought to study the impact of NB in *Cannabis* sp. growth, physiology, and yield. Four photoperiod treatments were compared: long day (T1: 18 h light/6 h dark), night break treatments with 12 h light photoperiods supplemented with 1 h light pulse (T2, at ZT+17)) or with four light pulses of 15 minutes each (T3: ZT+15 to ZT+18) and short-day treatment (T4: 12h light /12h dark). The appearance of pistils and reduction of internodes in apical segments were observed after 20 days only in plants subjected to short day (T4), while all four treatments displayed a similar physiological behavior. Regarding yield, plants with supplemental light presented higher biomass accumulation than the short-day treatment. Cannabinoid accumulation profiles showed distinct trends: CBD accumulation showed no differences, while THC accumulation was lower in T3 and T4 compared to T1 and T2. These results suggest that night interruption in non-psychoactive plants is as effective as extending the light phase in preventing inflorescence formation, without generating stress or reductions in yield.

## Introduction

The genus *Cannabis sp.* L (hereafter referred to as cannabis) belongs to the family Cannabaceae and groups dioecious or monoecious herbaceous plants native to Eurasia (McPartland, 2018; Q. Zhang *et al.*, 2018). Cannabis is traditional in this region as a source of fiber and food, and dates back around 6,000 years (McPartland *et al.*, 2018); however, today it is known and cultivated around the world primarily for the accumulation of secondary metabolites known as cannabinoids. These metabolites are synthesized in the trichomes of female inflorescences (Livingston *et al.*, 2020; Punja *et al.*, 2023). The harvesting of flower clusters for the extraction of ι19-tetrahydrocannabinol (THC), cannabidiol (CBD) and cannabigerol (CBG) among others, is widely used in the cosmetic and pharmaceutical industry for its analgesic, anti-inflammatory, anxiolytic, antispasmodic and antipsychotic properties (Andre *et al.*, 2016).

The most studied commercial trait of the plant is the accumulation of cannabinoids in female inflorescences (Aliferis & Bernard-Perron, 2020; Stack *et al.*, 2021; Dang *et al.*, 2022). One of the particularities of cannabis lies in the photoperiod-dependent formation of inflorescences, but not floral induction (Spitzer-Rimon *et al.*, 2019). Photoperiod refers to the length of the light phase in the day (24 hours) and is a strong influence on the circadian cycle (Creux & Harmer, 2019; de Montaigu *et al.*, 2010). According to Spitzer-Rimon *et al.* (2019), cannabis plants, even when exposed to prolonged photoperiods, mark the beginning of the reproductive stage with the appearance of solitary bracts and flower primordia; that is, cannabis plants generate solitary flowers as early as three weeks after rooting, but the development of inflorescences (flower clusters) is strongly determined by day length.

Most cannabis varieties described require long photoperiods (greater than 14h) to generate biomass and elongate branches, and short photoperiods to induce flower clusters (Hall *et al.*, 2012; M. Zhang *et al.*, 2021). For production in areas with short photoperiods, it is necessary to artificially lengthen the photoperiod to generate sufficient biomass to maintain the yield of clusters and cannabinoids per plant (Dang *et al.*, 2022). Previous studies have shown divergence in biomass and cannabinoid accumulation associated with photoperiod in both vegetative and inflorescence generation phases; plants exposed to long photoperiods for a short time have a higher biomass generation, while plants exposed to long photoperiods for a longer time have a high concentration of cannabinoids (Peterswald *et al.*, 2023). The accumulation of cannabinoids is also given by the time in which the harvesting of inflorescences is performed. It was previously reported that CBD accumulation is higher than that of ι19-THC under prolonged photoperiod conditions, even in strains with high THC contents. The electric energy used to artificially prolong the photoperiod means an impact of about 5% on production costs.

An alternative to the extension of the light phase is night disruption or Night Break (NB). This treatment has been widely used for the evaluation of photoperiodic control of growth and flowering and has been shown to be effective in inhibiting flowering under inductive photoperiods (Ishikawa *et al.*, 2005, Higuchi *et al.*, 2012, Cao *et al.*, 2016). In crops with short photoperiod-dependent flowering such as chrysanthemum (*Chrysanthemum morifolium*) and tomato (*Solanum lycopersicum*), plant flowering is suspended when the night phase is interrupted by light pulses, either blue, red or white (full spectrum) (Cao *et al.*, 2016; Nissim-Leví *et al.*, 2019). In cannabis, Whipker *et al.* (2019) concluded that day lengthening and the use of NB prevent the development of flower clusters. However, the influence of night break on plant architecture and cannabinoid production and the physiological consequences are unknown. In the present report we aimed to evaluate the responses of non-psychoactive cannabis plants to two types of night break. We evaluated physiological responses during the treatment phase, tracked the phenological progress of plants subjected to four different photoperiods, and generated data on biomass accumulation and cannabinoid concentrations at harvest. The results presented may be promising for reducing production costs and environmental impact of the crop.

## Materials and Methods

### Plant material

Plants of asexual origin of *Cannabis sp*, from the strain Stellita CBD, belonging to the gene bank of the company La Santa SAS, were used (https://www.lasanta.com/en/home-english/). The plants were acquired four weeks after cutting, already rooted. For the phenology and production experiment, the plants were planted in 20 L slabs, containing substrate composed of two granulometries of washed and buffered coconut husk: 30% of granulometry between 0.5 and 12 mm, and 70% of granulometry between 0.5 and 25 mm, according to the supplier’s technical data sheet (Saenz-Fety®). The planting density was 6 plants per m^2^. Fertilization of the plants during the whole experiment was done with NPK 12-5-40 (Van Iperen International) during the first four weeks, and 20-20-20 during the rest of the experiment; fertilization was done daily by fertigation from Monday to Saturday. Pruning management was carried out as follows: pruning of the apical shoot 2 weeks after sowing (WAS), and low pruning was carried out between the fourth and sixth WAS, and branches that did not have sufficient thickness (≤ 3 mm) were eliminated.

### Night Break treatments

Plants were placed under the same greenhouse in four areas completely isolated by black plastic, which prevented light contamination. 20W 6500 K bulbs with a photosynthetic photon flux density (PPFD) between 1758 and 2296 μmol.m^-2^.s^-1^ were used at a height of 2.10 m above the beds. Plants were grown with a photoperiod of 18 hours of light and 6 hours of darkness (18:6) for 30 days (adaptation period). Subsequently, the plants were distributed into four treatments (Fig. 1A): treatment 1 (T1) subjected plants to a commercial photoperiod of 18 h light and 6 hh darkness; treatment 2 (T2), plants were exposed to 12 h photoperiod plus night interruption treatment for one hour daily from 23 h to (18 h post dawn: ZT+17) 24 h (ZT+18); treatment 3 (T3), plants were grown under 12 h photoperiod and treated with four light pulses, each of 15 min with 45 min gaps between each other starting at 20 h 45 min (ZT+14); i.e., the first pulse was given from 20: 45 hours until 21 hours, the second, from 21:45 until 22 hours, the third pulse from 22:45 until 23 hours, and finally, from 23:45 hours until 00:00 hours; the fourth treatment (T4) was the short day photoperiod, of 12 light and 12 hours dark. The light treatments had a duration of 30 days.

**Figure 1.**
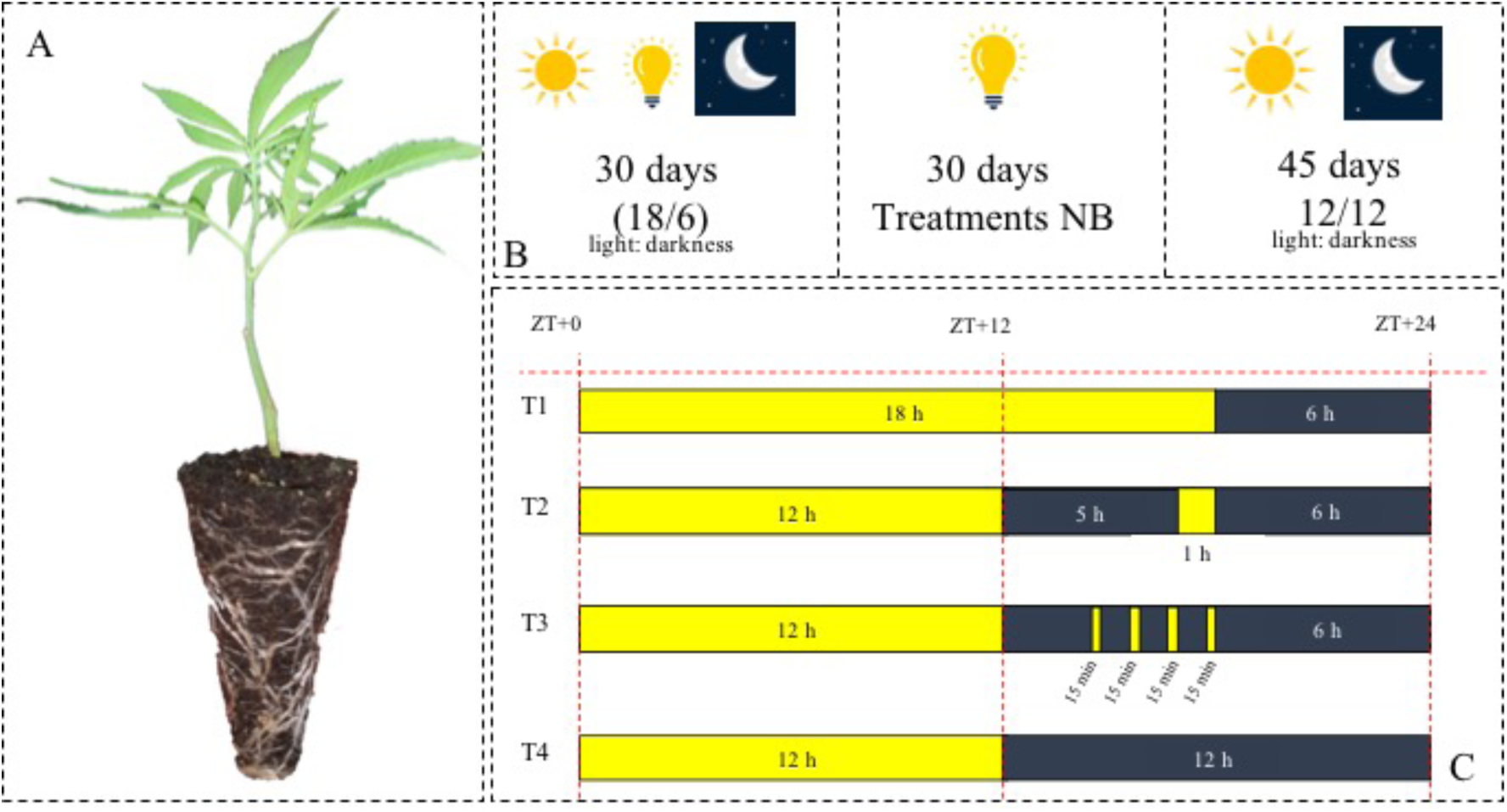
Experimental design. The culture cycle consisted of three stages: A. Cannabis sp. plant after 30 days of rooting, under non-inductive conditions (18/6). B. Cultivation stages of the experiment; 30 days for establishment of the plants under non-inductive greenhouse conditions; 30 days subjected to photoperiod treatments, and 45 days under 12:12 conditions (inflorescence induction).C. Photoperiod four treatments: T1 commercial photoperiod of 18 h light and 6 h darkness with 6 h of artificial light supplementation; T2 12 h of natural light plus 1 hour NB at ZT+17; T3 12 h of natural light and four pulses of 15 minutes, starting at ZT +15; and T4, control treatment 12 h of natural light and 12 h of darkness. Photographs by the author.

### Height, leaf area, internode size and floral advance curve

The phenological stages of *Cannabis sp.* plants subjected to light treatments (Night Break) were determined as follows (adaptation to the BBCH scale, 2nd Edition; Meier, 2001): the last vegetative stage (Main growth stage 4: vegetative propagation/starting, main shoot) was used for plants without pistils and marked as "49". Plants with pistils were identified with the code the first flowering stage (Principal growth stage 5: inflorescence emergence, main shoot). The presence of pistils in the thirds of the plant was used as a determinant of progress of floral development, and were scored according to the third where pistils were observed: in the lower third, they were identified with the number "52", middle third "54", middle and lower third "55", upper and middle third "56", and all thirds as "59".

In order to statistically analyze the phenological data, the above data were plotted as adjusted phenological stage (APS), converting the response variable into a response equivalent between 0 and 1, where 0 refers to the vegetative stage, and 1, to the most advanced flowering stage. An F1LDF1 design was used according to the taxonomy of Noguchi *et al.* (2012), using treatments (4 levels) as the between-subjects factor and time (6 levels) as the factor in repeated measures or longitudinal. The statistical model underlying this design can be described by independent random vectors Yik = (Yik1, . . ., Xikt)’, k = 1, ni, with marginal distributions Yiks ∼ Fis, i = 1, . . . . ., 4; s = 1, 11.

### Stomatal conductance and leaf temperature

For the analysis of stomatal aperture and leaf temperature, data were taken from the main leaflet of leaves in the middle third of the plant with the SC - 1 leaf porometer (Steady State Diffusion Porometer, Decagon Services, Washington, USA). Five plants per treatment were used, at two samplings time 5 and 20 Days After Treatment Induction (DATI) every 4 hours during 24 hours, starting at ZT+14 (8:00 pm).

### Chlorophyll a fluorescence

Chlorophyll a fluorescence data was taken with an unmodulated pulse fluorometer (Pocket PEA, Hansatech Instruments, UK) on leaves in the middle third, using 5 plants per treatment, at 5 and 20 DATI in a similar manner to the stomatal conductance data. Measurements were recorded at six different times throughout the day every 4 hours, starting at ZT+15 (8:00 pm). Initial fluorescence and peak fluorescence were measured to calculate the maximum quantum efficiency of photosystem II according to the equation noted in Supplementary Table 3 (Murchie & Lawson, 2013).

### Fresh and dry mass yield

Three treatments (T1, T2, T3) were harvested 105 days after sowing, while T4 was harvested 75 days after sowing to maintain the same time under conditions for inflorescence induction. The floral clusters produced per plant were harvested and subsequently weighed, discarding branches and leaves that did not contain trichomes. The fresh mass was taken from the harvest of the flowers of 8 plants for each of the four treatments, after 45 days of inductive photoperiod, cutting all the flowers that the plant had. The fresh mass was calculated with the average obtained from the 8 samples, using an Ohaus 3000g/0.1g balance.

The harvested flower, for all the analyses of all the treatments, was homogenized being this the initial weight (IW); after this, the flowers were taken to the drying oven (M.S.A. SAS) previously established at a constant 105 °C, where they remained for 90 minutes; after this time, the samples were weighed, and this value was taken as the final weight (FW). The percentage of dry matter was calculated according to the formula described in supplementary table 3.

### Analysis of cannabinoids

For the quantification of cannabinoids, 1 g of dry flower was taken from each of the 8 plants sampled for quantification in 3 replicates for each treatment. The quantification of cannabinoids (expressed in percentages) was done in the analytical laboratory KELAB (Bogota, Colombia). The concentration of cannabinoids in was measured by high performance liquid chromatography (HPLC). Method validation, along with repeatability, reproducibility and linearity was performed based on ICH HARMONISED TRIPARTITE GUIDELINE, provided by the European Medicine Agency (1995) and Somatek Ink. (2014) (https://somatek.com/wp-content/uploads/2014/06/sk140605h.pdf.

Five grams of dried inflorescence were sent to the analysis laboratory under room temperature conditions, to which the internal moisture was measured using a thermobalance. A subsample of 500 mg of dry flower was extracted to which a volume of 2 mL of diluent (H2O:MeOH) analytical grade was added in ratio 80:20 and vortexed for 2 minutes; an aliquot of the supernatant was taken to a volumetric balloon, "MS-stock solution". Again 200 mg were taken and 2 mL of the diluent were added, taking it to vortex again for 2 minutes at maximum revolutions, naming the sample as "stock solution extract-SSE".

Absorbance was measured on a UV detector at 228 nm on a 3 μm C18 column (Phenomenex, Torrance, CA, USA), 150 mm in length and 2.1 mm internal diameter, with a gradient flow (Table 1). The linearity curve was determined in the range of 0.1 and 200 mg/mL.

**Table 1.** Equipment and implements used in the quantification of cannabinoids by the KELAB Analytical of the KELAB Analytical laboratory.

| DEVICE | IMPLEMENT |
| --- | --- |
| **Column** | C18 |
| **Type of flow** | Gradient |
| **Detector** | UV 228 nm |
| **Diluent** | Methanol: Water |

| DEVICE | IMPLEMENT |
| --- | --- |
| Column | C18 |
| Type of flow | Gradient |
| Detector | UV 228 nm |
| Diluent | Methanol: Water |

The concentration (expressed in percentage) of cannabinoids (which was expressed on a dry basis), the concentration in (mg/mL), the conversion from concentration to percentage, the calculation for total CBD and total THC (CBDt and THCt), was performed according to the formulas in Supplementary Table 2.

The following were quantified: cannabidiolic acid (CBDA), cannabigerolic acid (CBGA), cannabidiol (CBD), cannabigerol (CBG) and ι19-tetrahydrocannabinol (THC) (in acid and decarboxylated form), cannabichromenic acid (CBCA), ι18-tetrahydrocannabinol and cannabinol (CBN), the latter three grouped as secondary cannabinoids.

### Experimental design and statistical analysis

A completely randomized design with four treatments was used. A multifactorial analysis of variance (ANOVA) with homogeneous variances and normal distribution was used. The effect of treatment, the effect of time and the effect of time on treatment were evaluated to analyze the effect of each test treatment. To evaluate floral development the analysis method used was the Longitudinal Nonparametric Analysis of Variance (ATS) (Noguchi *et al.* 2012).

A one-way ANOVA and Tukey’s multiple comparisons tests were used for the physiological variables, with a significance level of 0.05. Temperature, *gs* and Fv/Fm data were transformed by the Box-Cox method to comply with the assumptions of normality and homoscedasticity. Data processing and analysis was performed using R studio v. 1.4.1717.

## Results

### Plant height

In terms of the growth, cannabis plants subjected to different photoperiods did not show significant differences between treatments (Fig. 2A), with mean heights of 40 cm. However, it should be noted that leaf area did show significant differences between treatments. At the beginning of the treatments, the leaf area of the plants only showed significant differences between T3 and T4, and at 5 DATI we observed significant differences between T1 and T3. At 20 DATI there was a notable increase in leaf area in T2, while in the other treatments exhibited a stabilization in growth (Fig. 2B).

**Figure 2.**
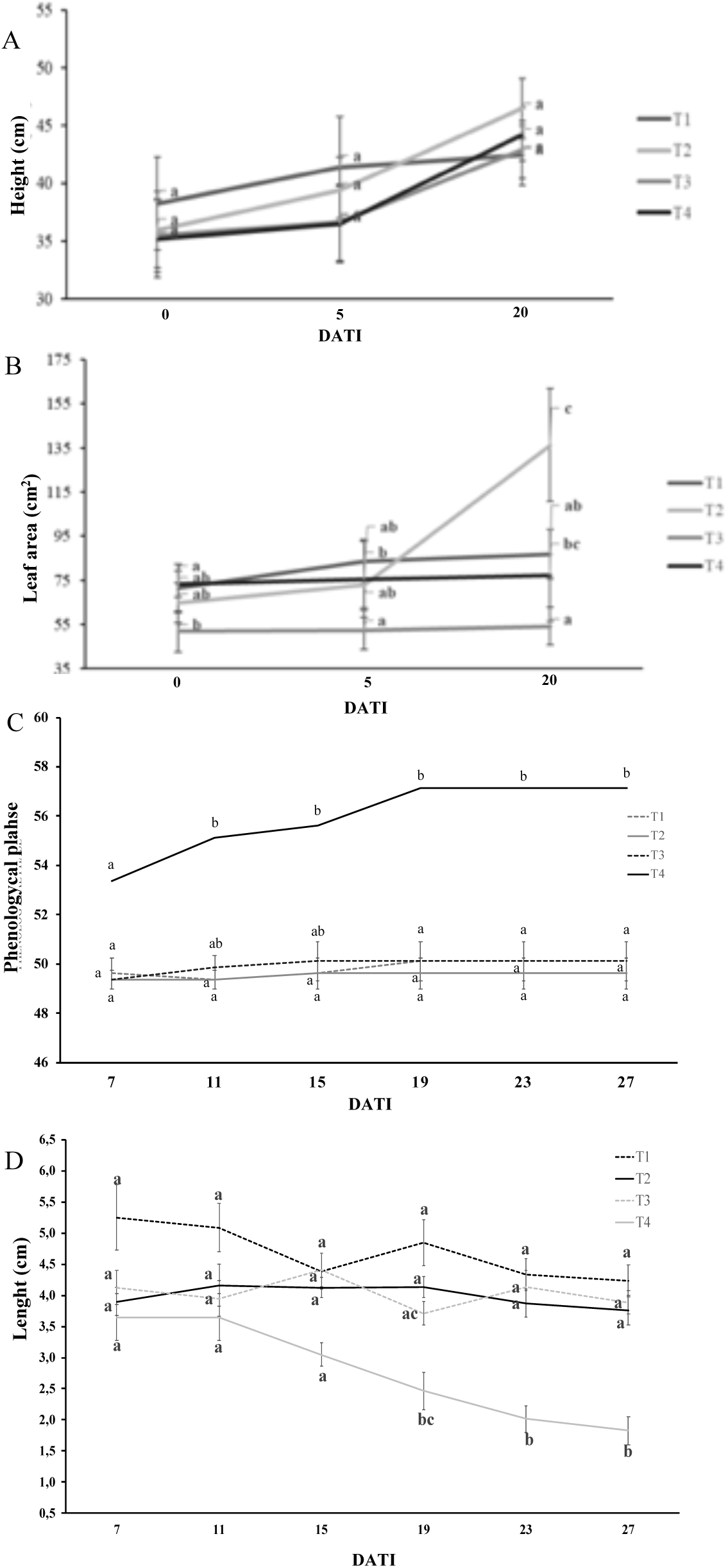
Morphological and developmental parameters in Cannabis sp. plants comparing four photoperiod treatments at three times: at the beginning of the treatments, 5 days after the beginning of treatments (DATI) and at 20 DATI, in 4 light treatments: T1 (18:6), T2 (NB with 1 h white light pulse), T3 (NB with 15 min light pulse, every hour for 4 h) and T4 (Control 12:12). (A) Plant height (n=8). (B) Leaf area (n=4). (C) Phenological advance according to the <u>Cannabis sp.</u> BBCH scale). (D) Internode length from the top shoot (n=8). Values presented are means for each treatment. Different letters above the bars represent significant differences between treatments by Tukey’s HSD test at α = 0.05.

### Phenology and BBCH scale

Plants in vegetative stage (49) were observed before day seven after treatment induction. Treatment 4 presented at 7 DATI the first flowers with visible pistils characteristic of stage 52, while the other three treatments remained in vegetative stage (Figure 2C) with the constant development of young vegetative buds. Despite this, no significant differences were found between the treatments. Treatment 4 progressed with time, until 100% of the plants were recorded at stage 59 in day 27 after treatments, characterized by the complete emergence of inflorescences, a step prior to stage 60 (initiation of flower formation). The appearance of reproductive structures was observed at 15 DATI for T4, but pistil development was observed in solitary flowers of all treatments, mainly in the lower and middle thirds independent of photoperiod.

In order to compare the results obtained between treatments, an additional step was performed to parameterize the categorical data obtained on the BBCH scale and thus, convert the response variable into an equivalent between 0 and 1 called relative effect (RE). The analysis shows that there are differences after 19 DATI between the three treatments with supplemental light and T4 (Supplementary fig. 1). Furthermore, the statistical analysis did not show any interaction of time with treatments (p=0.4438431) and no effect of time on plants under NB (p=0.2986930); with this, it is possible to interpret that the main effects of the study are based on the clear and strong effect of treatments on plants (p=1.758467e-42).

### Internode sizes

Measurement of the last three internodes showed differences among treatments at the start of the photoperiod treatments, although not significant. At 7 DATI, treatment 1 showed internodes averaging between 5 and 5.5 cm while the other treatments exhibited internodes 3.6 - 4.2 cm long (Fig. 2D); at 19 DATI, significant differences were observed comparing T4 with T1 and T2, while T3 and T4 were not significantly different. From this point until day 27, T4 significantly reduced internode lengths of the plants compared to the other treatments.

### Stomatal conductance and leaf temperature

The data from the measurement of stomatal conductance showed two similar patterns of behavior in the two sampling moments in the four treatments, with maximums at 12 noon and minimums at night, between ZT +14 and ZT +22. Although no significant differences were observed at any point at 5 DATI, we did observe a change at 20 DATI where significant differences in stomatal conductance maxima could be seen, with T3 showing the highest values and T4 the lowest values; for both cases, at ZT+6. A modulation of stomatal conductance was detected between sampling days, with higher gs values at 5 DATI, while at 20 DATI a reduction by almost half was detected at all hours sampled (Fig. 3A, 3C).

**Figure 3.**
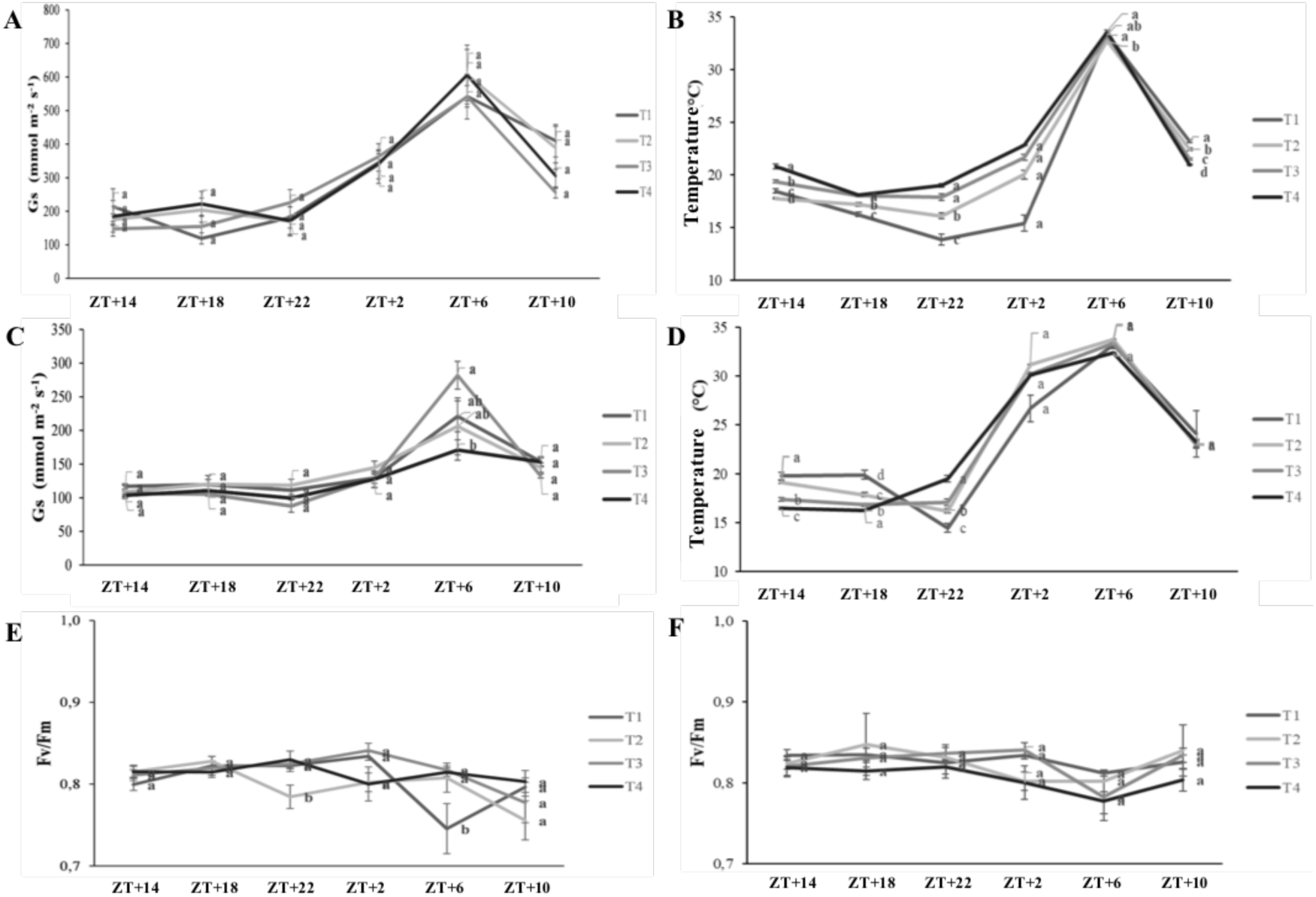
Physiological variables in Cannabis sp. plants comparing four photoperiod treatments: T1:18:6), T2: NB with white light pulse 1 hour, T3: NB with light pulses 15 min, every hour for 4 hours and T4: Control 12:12. A and B: stomatal conductance (gs) and leaf temperature taken during 24 h in leaves of Cannabis sp. plants taken at 5 DATI, respectively. C and D: stomatal conductance and leaf temperature taken for 24 h at 20 DATI, respectively. E and F: Maximum quantum efficiency of PSII photochemical activity (Fv/Fm) measured for 24 h taken at 5 and 20 DATI. Sampling time is expressed in zeitgeber time (ZT+0= dawn). Different letters indicate significant differences determined by Tukey test (HSD) at a level of P ≤ 0.05 (n=4).

Leaf temperature of plants subjected to the different photoperiods ranged from 15°C to 34°C. Significant differences between all treatments were evident between the evening and the morning. At 5 DATI, we found that plants showed a peak in temperature at ZT+6, with temperatures around 34°C, while the lowest temperatures were found at ZT+22 for T1, at which point significant differences were found against T3 and T4; statistically significant differences were also found for T2 compared with T3 and T4 (Fig. 3B). Significant differences were detected between T1 and T4 at almost all sampling points, with T4 registering higher temperatures. Treatments 2 and 3 showed more differences with T1.

At 20 DATI, a similar daily trend was detected for all treatments, with temperatures ranging from 15°C to 34°C, but peak temperatures (above 30°C) were detected at ZT+2 and ZT+6. Significant differences were observed between ZT+14 and ZT+18; T1 and T2 displayed the highest foliar temperature, while T3 and T4 exhibited the lowest values. At ZT +22 the trend reversed.

### Chlorophyll a fluorescence

Regarding the maximum quantum efficiency of PSII, values close to 0.8 were detected for all treatments, in general without significant differences at 5 DATI (Fig. 3E) and 20 DATI (Fig. 3F). At 5 DATI, significant reductions were evident in T2 at ZT+22 where a significant difference was evident against the other treatments; after this, we saw a partial increase until ZT+6 and again decreases at dusk, but no significant differences were detected. At 20 DATI the treatments showed no significant differences at any sampling time (Fig. 3F).

### Flower biomass

Significant differences in fresh mass were observed between treatments 4 and treatments 1 and 2 (Fig. 4A). Treatment 4, with 37.25 g/plant, had the lowest fresh flower weight yield, but it was not significantly different with respect to T3 (45.25 g/plant), while treatments 1 and 2 yielded 47.37 and 48.12 g/plant, respectively. In terms of flower dry mass, a significant decrease in dry matter content was detected for T4 with a total dry mass of 7.45 g, which was significantly different from the other treatments, which reached 13.81, 15.67 and 14.41 g for T1, T2 and T3 respectively (Fig. 4B), i.e., T2 showed a reduction of 47.54% with respect to the treatment with the highest production (T2). For treatment 2, 33.08% of the fresh sample is dry mass, followed by treatment 3, with 31.85%. In the case of treatment 4, the percentage of dry matter is 20%.

**Figure 4.**
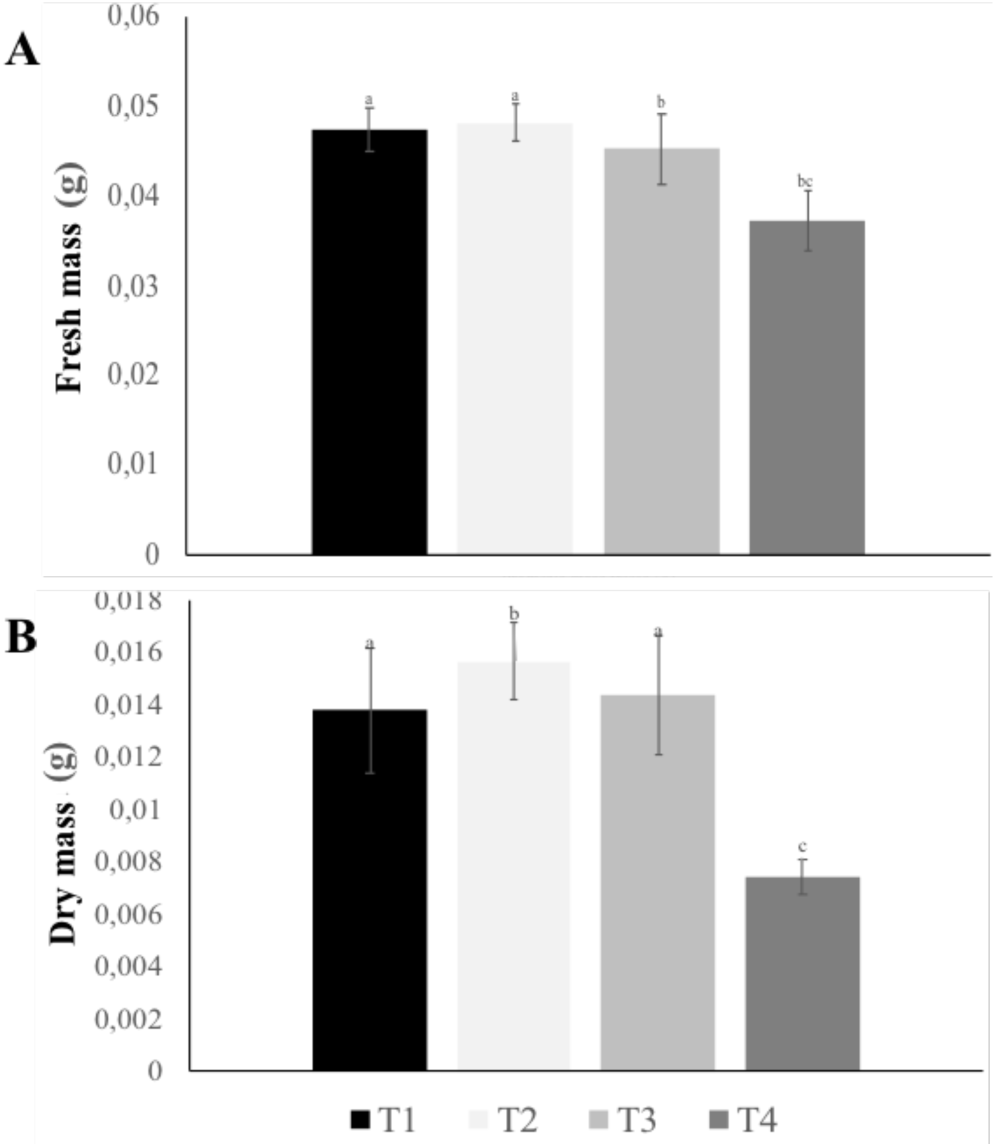
Yield of Cannabis sp. plants subjected to photoperiod treatments in four different treatments: T1 (18:6), T2 (NB with 1 h white light pulse), T3 (NB with 15 min light pulse, every hour for 4 h) and T4 (Control 12:12). (A) Fresh mass measured from harvested cannabis flowers. Dry mass of harvested cannabis flowers per treatment. Different letters above the error bars represent significant differences between treatments by Tukey’s HSD test at α = 0.05 (n=8).

### Cannabinoid content

The concentration of cannabidiol (CBD), cannabidiolic acid (CBDA) and total CBD (CBDt; Supplementary Table 1) was measured. The results of this experiment showed that CBD concentration (Fig. 5A) was significantly different in T2 with a concentration of 5.45 (% w/w) against the other three treatments; T1 was also significantly different (3.56% w/w) against T3 and T4, which yielded a concentration (%w/w) of 2.06 and 1.05, respectively. Treatments 3 and 4 showed no significant differences between them. CBDA concentration showed similarities between treatments 1 and 3, which yielded 3.98 and 4.71%, respectively. For treatment 2, the dried flower samples yielded the lowest CBDA concentration (0.98% w/w), while T4 obtained a CBDA concentration higher than 5% w/w (Fig. 5A). CBDt concentration had no significant differences for any treatment; however, T1 accumulated the highest concentration with 7.05% w/w, and the treatment with the lowest concentration was T4 with 6.04% w/w.

**Figure 5.**
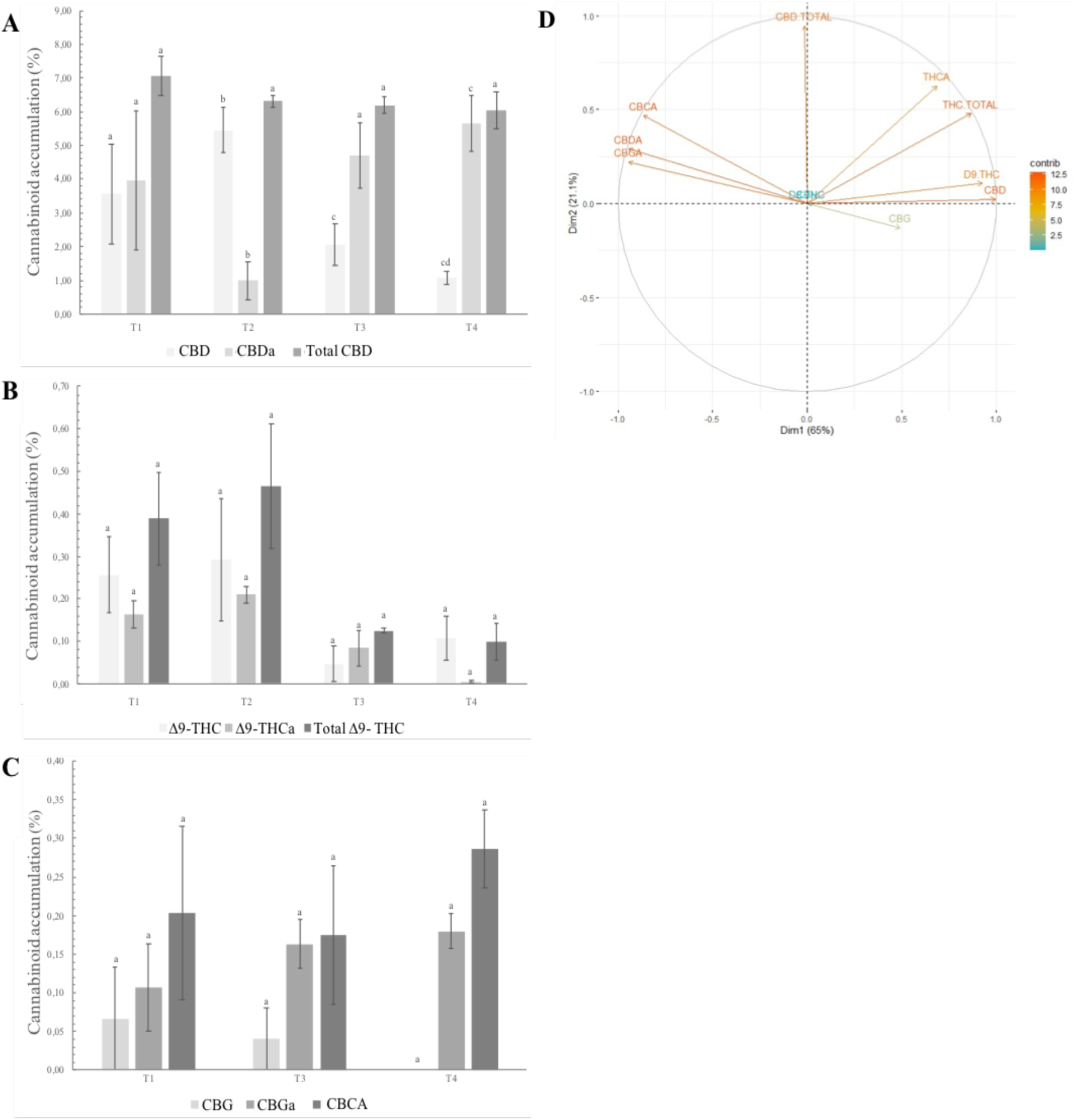
Cannabinoid accumulation in cannabis plants (% dry weight). subjected to four photoperiod treatments. (A) Concentration of cannabidiolic acid (CBDA), cannabidiol (CBD) and total CBD (CBDt) present in flowers (B) Concentration of Δ9-tetrahydrocannabinolic acid (THCA), Δ9-tetrahydrocannabinol (THC) and total THC (THCt) in flowers of cannabis plants (B) Concentration of cannabigerolic acid (CBGA), cannabigerol (CBG) and cannabichromenic acid (CBCA) in flowers. Different letters above the error bars represent significant differences between treatments by Tukey’s HSD test at α = 0.05 (n=3). (D) Biplot showing the contribution of each cannabinoid detected by HPLC to the total variability of the data obtained in plants subjected to photoperiod treatments.

The results for tetrahydrocannabinol (Δ9-THC) and tetrahydrocannabinolic acid (THCA) concentration remained below 1% and showed no significant differences between treatments (Fig. 5B). In the case of total tetrahydrocannabinol concentration (THCt), there were significant differences between T2 and T4, accumulating 0.47 and 0.10, respectively, but not between the other treatments (T1 = 0.39% and T3 = 0.12). It should be clarified that the chemotype used for the plants is non-psychoactive, whose values according to the supplier, do not exceed 1% of Δ9-THC.

The accumulation of secondary cannabinoids such as cannabigerol (CBG), cannabigerolic acid (CBGA) and cannabichromenic acid (CBCA) was also analyzed. The chromatography results indicate that T2 did not accumulate any of these cannabinoids; on the other hand, the plants from the other three treatments, although not significantly different from each other, accumulated low percentages. In no case was the concentration greater than 0.3% w/w (Fig. 5C).

The contribution of principal components was plotted, which resulted in a contribution of more than 80% of the variation of the data and was visualized by means of a biplot as (Figure 5D). We found that the presence of cannabinoids such as THC and CBD resulting in our analysis, contribute significantly to explain the variation of our results (>10%) in the experiment. A opposite correlation was observed between CBGA, CBDA and CBGA, versus THCA, THC and total THC (Fig. 5D).

## Discussion

One of the major costs in the production of cannabis for medicinal or industrial purposes derives from the prolonged use of electricity, which is about 5.8% of the total costs for the companies (Dang *et al.*, 2022). Day extension using artificial light prevents the development of flower clusters and prolongs the vegetative phase for the formation of productive branches. The present study shows that NB prevents inflorescence formation without affecting development by reducing lighting times.

It was observed that plants exposed to the evaluated photoperiods did not show significant differences in height. Contrary to what was previously reported by Whipker and collaborators (2019), plants under light treatments presented the same height as plants during the experiment. It should be noted that the previous report was made in incubators with artificial light, while our conditions were greenhouse conditions with about 12 h of sunlight. However, an effect on the leaf area of the plants was observed; the plants with the greatest leaf area were from T2, which, in turn, had a greater height (Fig. 2A-B). These differences in area may also be due to differences in hyponastic movements and growth under the photoperiods evaluated; The four photoperiods impose different light requirements for plants which might be traduced into variation in leaf movements, evident at our sampling time (Apelt *et al.*, 2017; Dornbusch *et al.*, 2014).

Indicative of the onset of inflorescence formation is the shortening of internodes (Fig. 2), the appearance of floral bract structures, increased trichome density, and marked reduction in leaf size (Spitzer-Rimon *et al.*, 2019; Sutton *et al.*, 2023). At the beginning of the experiment all plants displayed solitary flowers, in agreement with that reported by Spitzer-Rimon *et al.* (2019). In the experiment we found significant differences in internode size of plants evaluated up to 15 DATI with internode shortening only for T4, which corresponds with generation of racemes (Hesami *et al.*, 2023). The shortening of internodes was accompanied by sighting of the first apical flower buds, their clustering, the change of leaf arrangement from opposite to alternate, and decrease in petiole length (Mediavilla *et al.*, 1998; Mishchenko *et al.*, 2017). In addition, under an advanced flowering stage, flowers take the shape of smaller spikelets and have no petiole (Strzelczyk *et al.* 2022; Gill *et al.*, 2022; Horne, 2020; Nelson 2000; Small *et al.*, 2003; Srivastava and Yadav 2013; Vuerich *et al.*, 2019) which gives way to raceme formation. In conclusion, NB treatments generate an effect on plants similar to the currently used day length extension (18 hours of light).

On the other hand, plant physiology can be altered by abiotic events, such as photoperiod, light intensity, and light quality (Creux & Harmer, 2019; Poorter *et al.*, 2019; Slattery *et al.*, 2018). Monitoring of stomatal conductance and leaf temperature conducted for 24 h at 5 and 20 DATI detected diurnal fluctuations related more to environmental conditions than to photoperiod treatments (Fig. 3A, C), with maxima around midday (ZT+5 - ZT+7) and coinciding with the highest temperatures (Gill *et al.*, 2022; McAusland *et al.*, 2021). In addition, we observed significant changes in stomatal conductance at 5 and 20 DATI, possibly associated with the growth and maturity of the leaves evaluated; it was previously reported in *Pyrus serotina* that fully expanded young leaves tend to have high gs and as maturity advances gs is reduced, (Xie & Luo, 2003). On the other hand, higher foliar temperatures were detected at 20 DATI, which could mean higher environmental temperatures and reduced stomatal conductance.

Measurement of physiological parameters indicative of abiotic stress such as PSII quantum efficiency (Fv/Fm) was performed. The measurement of fluorescence parameters allows to qualitatively and quantitatively analyze the absorption and utilization of light energy through PSII in relation to photosynthetic capacity in a non-destructive way (Mouget & Tremblin, 2002; Baker & Rosenqvst, 2004; Jimenez-Suancha *et al.*, 2015). The values are very close 0.8, which allows us to affirm that the photoperiod evaluated do not generate stressful states as previously observed (Rodríguez-Yzquierdo *et al.*, 2021). Based on the above, we can affirm that Night Break treatments do not alter physiological parameters that could associated them to stress response. To our knowledge, this research is the first in which physiological values such as stomatal conductance and plant temperature of *Cannabis sp.* are considered.

In terms of yield, T1 and T2 accumulated more dry mass (Fig. 4), with T2 having significantly higher values, followed by T1 and T3. This indicates that the night interruption with one hour of light is as efficient in producing dry mass as the long day photoperiod, as previously observed with other type of NB (Whipker *et al.*, 2020). Treatment 4 had a significant reduction given most likely by its early exposure to short photoperiod conditions that accelerated inflorescence development at the expense of new branches and raceme generation.

Cannabinoid production (percentage of accumulation per g dry matter) showed differences in the cannabinoid profile between treatments; however, there were no significant differences in accumulation (Fig. 5). The variety used in the report is a variety characterized as non-psychoactive according to the Colombian registry and its producers (Hidalgo, *pers. comm.*). The concentration of total CBD per g of dry flower remained stable, suggesting that photoperiod does not affect the accumulation of this cannabinoid. However, the treatments do show some differences in CBDA and CBD content, which could be associated with abiotic factors such as temperature, light, air or heat that could have generated decarboxylation of CBDA (Chandra *et al.*, 2013; Citti *et al.*, 2018; Livingston *et al.*, 2020).

On the other hand, variation in the kinetics of cannabinoid accumulation over time between varieties (Yang *et al.*, 2020) and variation in the final accumulation of cannabinoids in plants exposed to different photoperiods (Peterswald *et al.*, 2023) has been reported. These phenomena could be related to the inflorescence maturation process (Naim-Feil *et al.*, 2023). The accumulation of cannabinoids in cannabis plants occurs progressively due to the maturation of the inflorescences and the progressive synthesis of the acidic forms of the cannabinoids in trichomes (Livingston *et al.*, 2020). According to Pacifico and collaborators (2007), exponential accumulation of cannabinoids begins after 60 days, until day 80-85 after rooting, which coincides with the time in which our harvest was carried out (45 DATI; 105 days after sowing for T1, T2 and T3, and 75 days after sowing for T4). During the maturation process of the inflorescence, the synthesis of THC, CBD and CBC is carried out from CBG (Gülck & Møller, 2020). A report by Burgel and collaborators (2020) shows accumulation of CBD and CBG in hemp plants, in early stages of the inflorescence and accumulation of THC in more advanced stages. Our results show two trends: one related to the accumulation of CBDA, CBCA and CBGA and another with the accumulation of THCA and CBD, which could be related to differences in inflorescence development stages between treatments. These findings allude to possible variations in the inflorescence maturation process influenced by the photoperiod of the previous phase.

In conclusion, the NB treatments tested on a non-psychoactive strain are as effective in preventing flower cluster induction as long day, without impacting physiology or dry flower yield. However, we did detect small differences in cannabinoid profiles suggesting a possible effect of treatment on subsequent inflorescence generation and cannabinoid accumulation. Future studies should focus on the kinetics of cannabinoid accumulation prior to and during flower cluster induction, and on the molecular factors associated with both night break effect and inflorescence maturation.

## Acknowledgments

The authors thank Professor Luz Marina Melgarejo and Ms. Paula Andrea Lozano, M. Sc. for the collaboration with the physiological experiments and analyses. The authors would also like to thank Mr. Javier Hidalgo for supplying the plants used in the experiments.

## Supplementary material

**Supplementary table 1.** Cannabinoid concentration of plants subjected to four photoperiod treatments.

| SAMPLE | CBD | CBDA | THCA | $\Delta^9$-THC | CBG | CBGA | CBCA | TOTAL<br>CBD | TOTAL<br>THC |
| --- | --- | --- | --- | --- | --- | --- | --- | --- | --- |
| T1 | 3,56 | 3,98 | 0,26 | 0,16 | 0,07 | 0,11 | 0,20 | 7,05 | 0,39 |
| T2 | 5,45 | 0,98 | 0,29 | 0,21 | 0,00 | 0,00 | 0,00 | 6,31 | 0,47 |
| T3 | 2,06 | 4,71 | 0,05 | 0,08 | 0,04 | 0,16 | 0,18 | 6,19 | 0,12 |
| T4 | 1,08 | 5,66 | 0,11 | 0,00 | 0,00 | 0,18 | 0,29 | 6,04 | 0,10 |

**Supplementary table 2.** Table of marginal mean comparisons by treatment of phenology analysis:

| Comparison | Hypothesis | P value | Adjusted P<br>valor |
| --- | --- | --- | --- |
| 18_6 y 12_12 | $F_{18_6} = F_{12_12}$ | 1.05e-32 | 6*1.05e-32 |
| 18_6 y NB_1h | $F_{18_6} = F_{nb1h}$ | 0.6695 | 1 |
| 18_6 y NB_15m | $F_{18_6} = F_{nb15m}$ | 0.7791 | 1 |
| 12_12 y NB_1h | $F_{12_12} = F_{nb1h}$ | 1.21e-30 | 6*1.21e-30 |
| 12_12 y NB_15m | $F_{12_12} = F_{nb15m}$ | 3.99e-30 | 6*3.99e-30 |
| NB_1h y<br>NB_15m | $F_{nb1h} = F_{nb15m}$ | 0.5095 | 6*0.5095 |

**Supplementary table 3.**
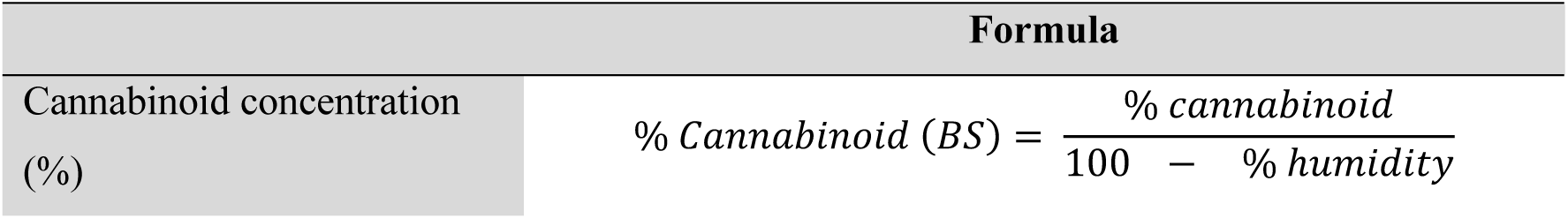
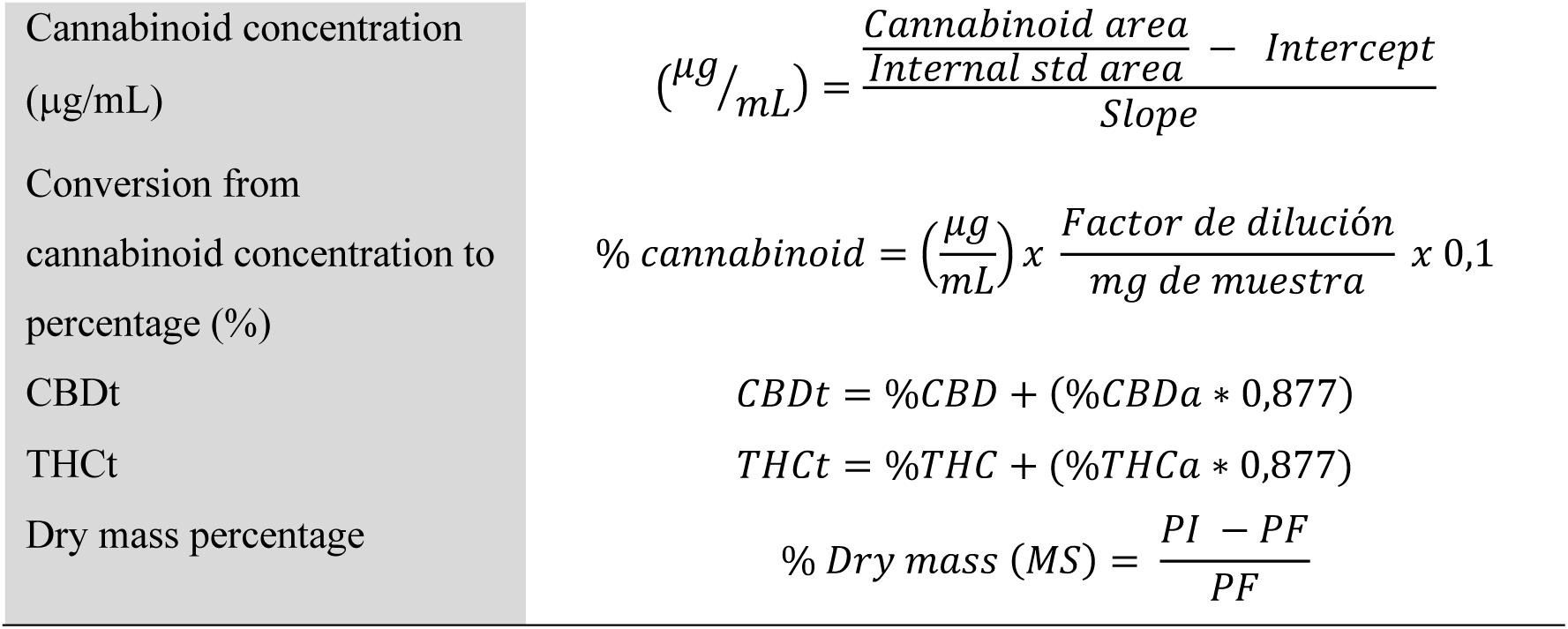
Formulas used in the experiments.

**Supplementary figure 1.**
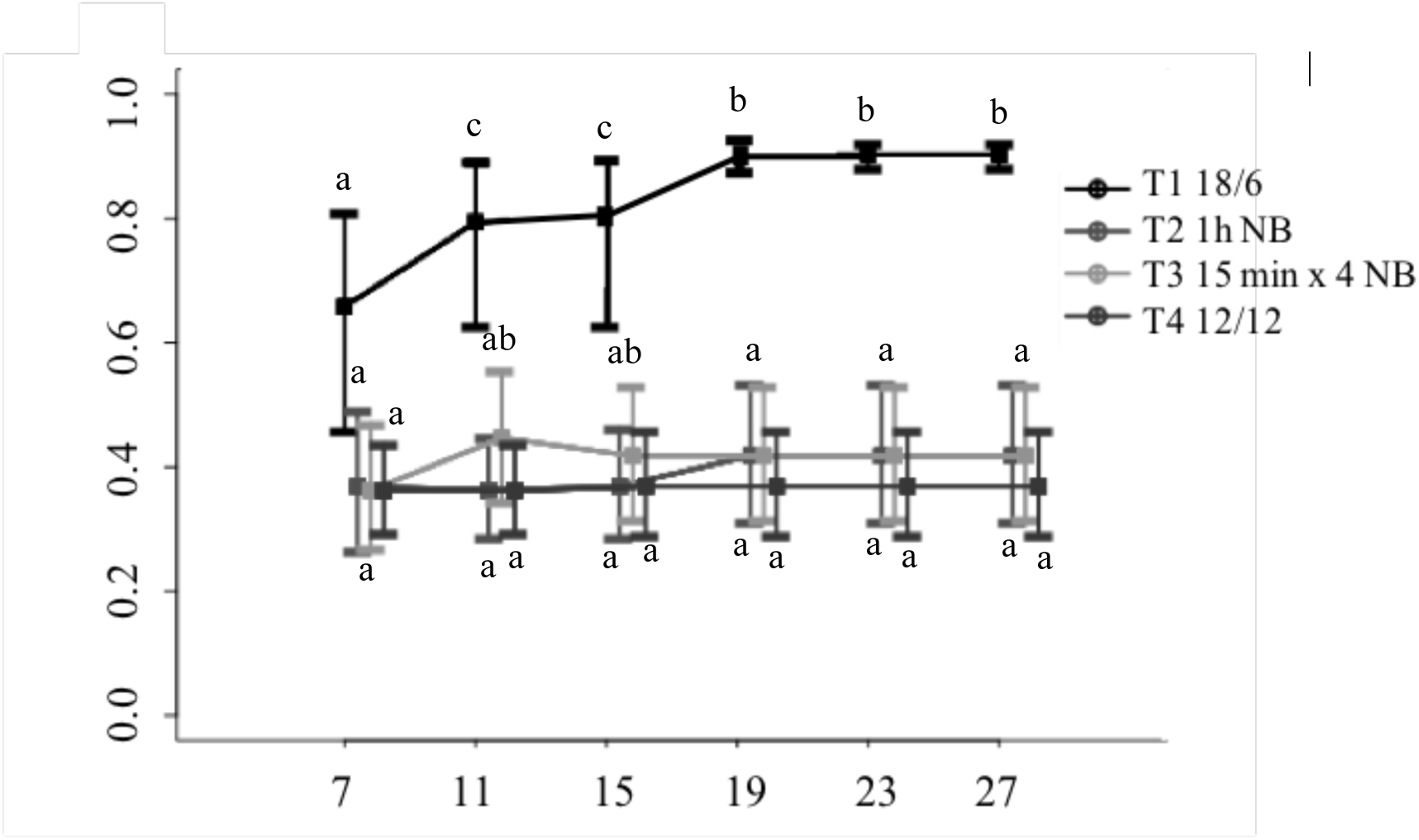
Adjusted phenological advance (BBCH scale). Phenological data were plotted as adjusted phenological stage (APS), converting the response variable into a response equivalent between 0 and 1, where 0 refers to the vegetative stage, and 1, to the most advanced flowering stage.

